# Single-cell dissection of cell hierarchies in urine-derived stem cells – application to chondrogenesis and general scalability

**DOI:** 10.1101/2025.09.26.678723

**Authors:** Alexander Schulz, Emily Brockmann, Miriam Zentgraf, Andreas S. Baur, Steffen Uebe, Arif B. Ekici, Mark Dedden, Sebastian Zundler, Christian T. Thiel

## Abstract

Developing of models of human cartilage and bone growth is essential for the study of growth disorders and to advance towards personalized therapeutic interventions. The majority of *in vitro* strategies depend on the use of invasively obtained mesenchymal stem cells (MSCs) or the laborious generation of induced pluripotent stem cells (iPSCs). We have now established urine-derived stem cells (USCs) as a non-invasive stem cell source capable of osteogenic and robust chondrogenic spheroid differentiation. Single-cell RNA sequencing of USCs revealed a hierarchy originating from parietal epithelial cells of the kidney, with a proliferative *TOP2A*⁺ subpopulation governed by *MYC* and *E2F4* regulatory networks. Pseudo-time analysis of chondrogenic USCs uncovered alternative chondrogenic differentiation trajectories with an *ALDH1A2*⁺ intermediate state and a *TIMP3*⁺-expressing chondrocyte-like subpopulation as the major endpoint, exhibiting cartilage-specific gene ontologies. In conclusion, a streamlined, xeno-free culture and differentiation protocol was developed, thereby establishing the basis for clinical-grade cell expansion and cartilage matrix formation. This positions USCs as a powerful tool for studying cartilage biology and a potential platform for development and use in regenerative therapies.

## Introduction

Developmental disorders of the skeleton have major implications for human health, but their underlying mechanisms remain only partially understood^1^. They manifest clinically as impaired longitudinal bone growth leading to short stature and affecting 2.3% of the general population^2^. Growth disorders can be attributed to various etiologies, including complex genetic syndromes^3-6^, endocrine defects^7-9^, skeletal dysplasias^10,11^, chronic diseases^12^, and idiopathic short stature^13,14^. At the cellular level, the pathology may originate from any zone of the growth plate (reserve, proliferative, or hypertrophic) or from the metaphysis and epiphysis^15,16^. Disruption of the function of the cells that comprise the cartilage (chondrocytes) and the composition of the extracellular matrix (ECM) have been shown to impair the organization of growth plates, leading to abnormal bone elongation^15^. Known genetic causes include genes encoding growth plate matrix proteins, such as *COL2A1* associated with spondyloepiphyseal dysplasia and hypochondrogenesis^17^, or *ACAN* resulting in premature growth plate closure^18^. Defects in the key regulators of cartilage cells such as Indian hedgehog, underlying brachydactyly type A1 and acrocapital femoral dysplasia^17^ are also well known. Within the aforementioned spectrum, skeletal dysplasias are of particular pertinence, with achondroplasia representing a prevalent form affecting more than 360,000 individuals worldwide^19^.

However, research on disorders of cartilage and bone formation is hampered by the limited access to primary human chondrogenic tissue, as it requires invasive procedures with potential harmful side-effects^20^. Consequently, the functional study of skeletal disorders and the development of patient-specific therapies is challenged by a paucity of suitable and broadly accessible cellular models. The majority of *in vitro* models used for studying chondrogenesis utilize either induced pluripotent stem cells (iPSCs) or mesenchymal stem cells (MSCs). MSCs, which possess the capacity to differentiate into cartilage, are obtained through an invasive procedure from tissue sources such as adipose tissue, bone marrow, or synovial fluid^21,22^. In contrast, iPSCs offer the benefit of a pluripotent and renewable source. However, these methods are expensive, time-consuming, and laborious, which limits their availability^23,24^. Moreover, their therapeutic value is further constrained by intrinsic tumorigenicity^25^. Consequently, both MSC and iPSC approaches exhibit significant practical limitations when modeling and treating genetic skeletal disorders, particularly in cases requiring repeated sampling, or in patient-specific studies.

Urine-derived stem cells (USCs) have emerged as a promising alternative to overcome these challenges. These can be obtained without the need for surgical procedures and easily propagated and mantained^26^. Despite increasing interest in their cartilage regenerative capabilities^27-29^, the application of USCs as a model system for cartilage and skeletal diseases has remained largely unexplored, particularly in the context of developmental skeletal disorders. In addition, the conventional culture of USCs, as initially identified in their original discovery^30^, is complex and reliant on animal-derived products, preventing potential use in cell-therapeutic approaches due to the inherent risks associated with the transfer of animal factors to patients^31^.

We here propose USCs as a platform for investigating human chondrogenesis and modeling of genetic skeletal disorders. Furthermore, a simple, xeno-free culture and chondrogenic differentiation method for USCs is presented, which has the potential to pave the way for personalized, non-surgical medicine and cartilage regeneration.

## Results

### USCs are parietal epithelial cells derived from *MYC*- and *E2F4*-driven stem cells

We performed scRNA-Seq to identify and characterize USCs. After standard Seurat^32^ QC procedures and clustering of 19,024 cells, nine transcriptionally distinct clusters were identified (UMAP, Fig. 1A, B, Supplementary Fig. 1A–E). We confirmed the kidney origin of USCs by comparing with established urinary and renal genelzsets (CellMarker 2.0^33^) (Fig.□1C). Anchor-based data transfer from a published kidney atlas^34^ revealed a parietal epithelial cell (PEC) identity of USCs. The expression of PEC-specific markers was observed to be highest in cluster 4, which also demonstrated the highest PEC-marker expression and mapping confidence (Fig. 1C–F).

**Figure 1:**
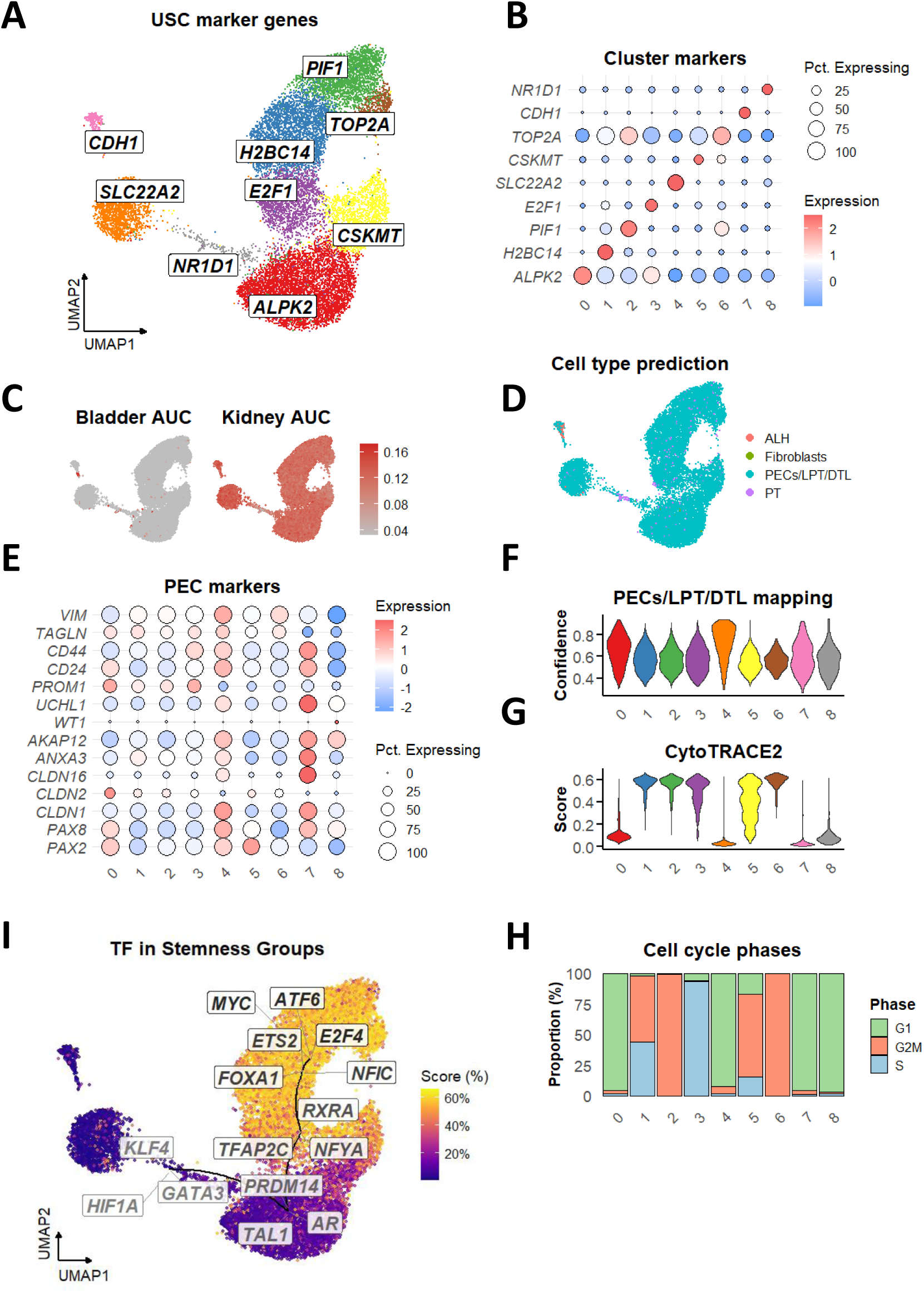
Transcriptional identity and stemness architecture of USCs. A: Uniform Manifold Approximation and Projection (UMAP) of USCs with biological top markers annotated per cluster. B: Dotplot of biological top marker expression across cluster. C: AUC-based enrichment scores of bladder and kidney-specific expression profiles projected onto UMAP space. D: UMAP of predicted cell types from label transfer of kidney reference cell types. (Cells of ascending loop of Henle = ALH, of the parietal epithelium = PECs, of late proximal tubule = LPT, of distal thin limb = DTL, of proximal tubule = PT). E: Dotplot showcasing expression of PEC markers among clusters. F: The mapping confidence of individual clusters to the PECs/LPT/DTL of reference. G: CytoTRACE2 scoring of individual clusters. H: Cell cycle phase proportions of individual clusters. I: Top 3 TFs per stemness group overlaid onto CytoTRACE2 score (Score) based UMAP in a percentage-based color gradient manner. (darker boxes = more stemness and vice versa.)

However, given that only two clusters showed a PEC marker gene expression signature, and USCs were previously reported to have multipotent differentiation capabilities^35^, we investigated the drivers of stemness in these cells. The application of CytoTRACE2^36^-based stemness scoring revealed a negative correlation between PEC identity and high potency levels, particularly in the *TOP2A*⁺ cluster in G₂/M phase. Analysis of transcription factor activity using VIPER, further identified *E2F4* and *MYC* as master regulators of the stem-like state, whereas *KLF4* activity marked quiescent PEC-like clusters (Fig. 1G–I, Supplementary Fig. 2B).

### USCs show osteogenic and chondrogenic differentiation potential

In order to develop cellular models for growth disorders, we analyzed and confirmed the differentiation potential of USCs towards osteogenic and chondrogenic differentiation. CytoTRACE2 potency UMAP revealed that a considerable proportion of cells showing a multipotency expression profile (Fig. 2A), suggestive of the capacity to generate multiple cell types. Thus, USCs were cultured in differentiation media for 2 weeks, followed by staining of osteogenic-induced cells positive for calcium deposits for their osteogenic potential. Chondrogenic cultures were positive for glycosaminoglycans (GAG) and formed chondrogenic aggregates, resembling normal chondrogenic differentiation^37^ (Fig. 2B).

**Figure 2:**
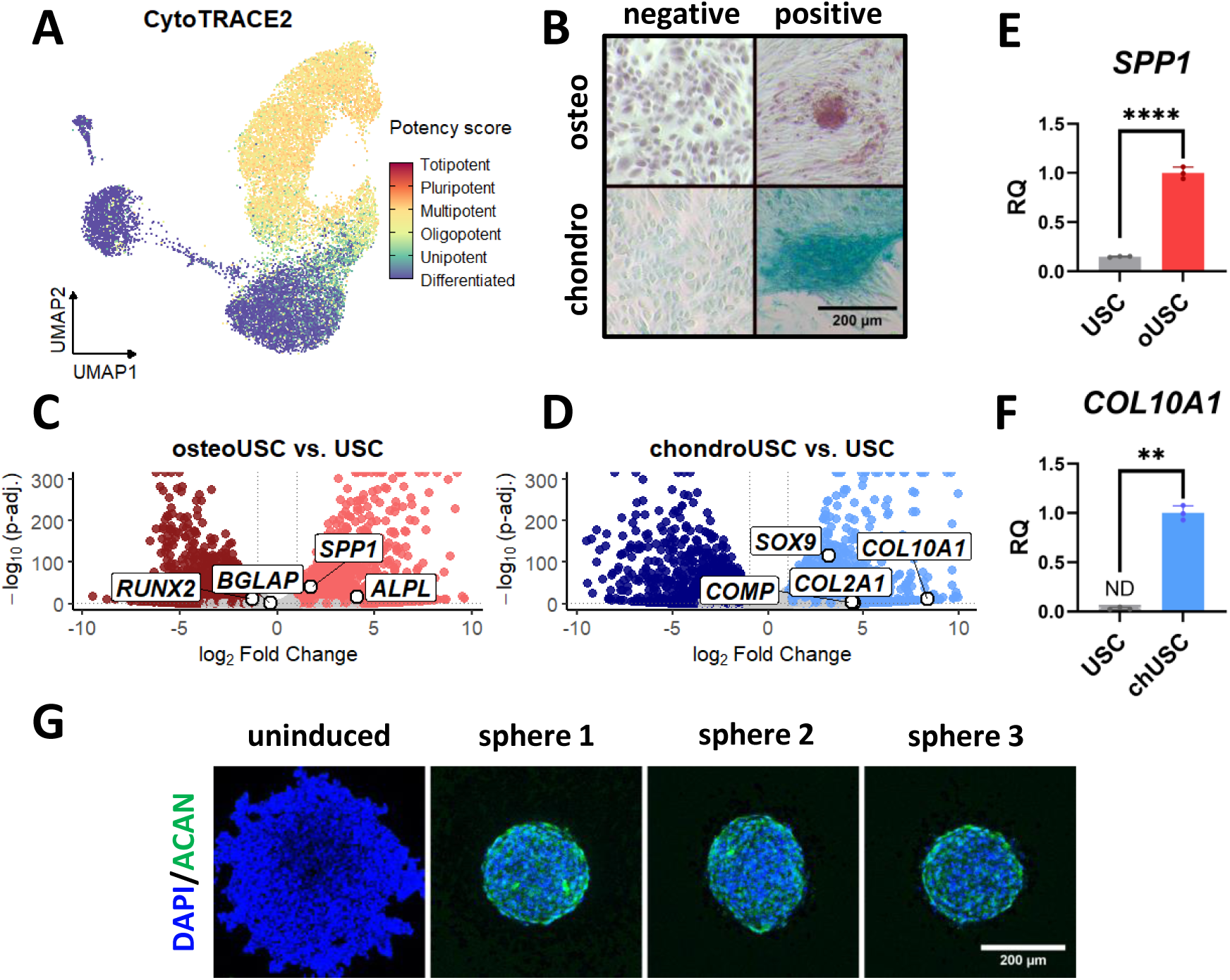
Functional differentiation capacity of USCs. A: CytoTRACE2 potency UMAP depicting differentiation capacities across cells. B: GAC and calcium staining of uninduced (negative) or induced (positive) cells. (osteo= osteogenic differentiation stained with alcian blue, chondro= chondrogenic differentiation stained with alizarin red). C/D: Volcanoplots of osteogenic/chondrogenic differentiation, with genes expressed canonically in osteoblasts/chondrocytes labbelled (genes highlighted: adjusted p-values < 0.05, log2 fold change > ± 1). In osteogenic USCs 2/4 genes are positively upregulated, in chondrogenic 4/4. E: *SPP1* expression levels in undifferentiated (USC) and osteogenic (oUSC) cells as measured by RT-qPCR. RQ undifferentiated = 0.149 ± 0.006, differentiated = 1.00 ± 0.0.06, n = 3, p < 0.0001, Welch’s t-test. F: RQ values of *COL10A1* of USCs (USC) and chondrogenic (chUSC) cells. Bar with undetermined CT-values or exceeding 40 cycles, was marked (=ND). *COL10A1*, RQ undifferentiated = 0.030 ± 0.013, differentiated = 1.00 ± 0.07, n = 3, p < 0.01, Welch’s t-test. G= Immunofluorescence staining of uninduced (negative), or chondrogenically induced spheroids (sphere 1-3). (Scale bar represents 200 µm).

Subsequently, we sought to ascertain whether these cells also express bona fide markers for osteoblast and chondrocytes. Bulk RNA-Seq and reverse transcription quantitative polymerase chain reaction (RT-qPCR) revealed expressed gene patterns of chondrocyte and osteoblast lineage specification (Fig. 2C-F, Supplementary Fig. 2C). Chondrogenic cultures exhibited a more pronounced upregulation of lineage-specific genes (Fig. 2D) in comparison to osteogenic cultures (Fig. 2C), (osteogenic: 2/4, chondrogenic 4/4 canonical genes), while both conditions increased expression of genes associated with terminal differentiation, as confirmed by qPCR (Fig. 2E, F). In order to confirm these results from monolayer cultures in 3D, we generated 3-week chondrospheres through cell-aggregation in low attachment plates. After differentiation, we detected strong aggrecan immunofluorescence (Fig. 2G, Supplementary Video) as a marker of articular cartilage, confirming that USCs form bona fide cartilage matrix in 3D culture. Taken together, our data demonstrates that USCs can differentiate along both osteogenic and especially chondrogenic pathways.

### Chondrogenic differentiation of USCs at single cell resolution

To unravel how urine-derived stem cells (USCs) acquire a chondrogenic identity at the cellular level, we analyzed single-cell RNA sequencing data from undifferentiated and chondrogenically differentiated monolayer cultures (Fig. 3). This approach allowed us to trace individual transcriptional trajectories and determine whether USC differentiation recapitulates key steps of endochondral ossification, the natural process driving cartilage formation during skeletal development. We first integrated 10,325 cells from the chondrogenic stage with a healthy cartilage reference atlas (Supplementary Fig. 1F–J). This enabled us to compare USC-derived cells to genuine chondrocytes and detect subpopulations that resemble the cartilage lineage (Fig. 3A). This finding indicates that USCs undergo a transcriptional maturation process, mirroring early-to-late chondrogenic commitment.

**Figure 3:**
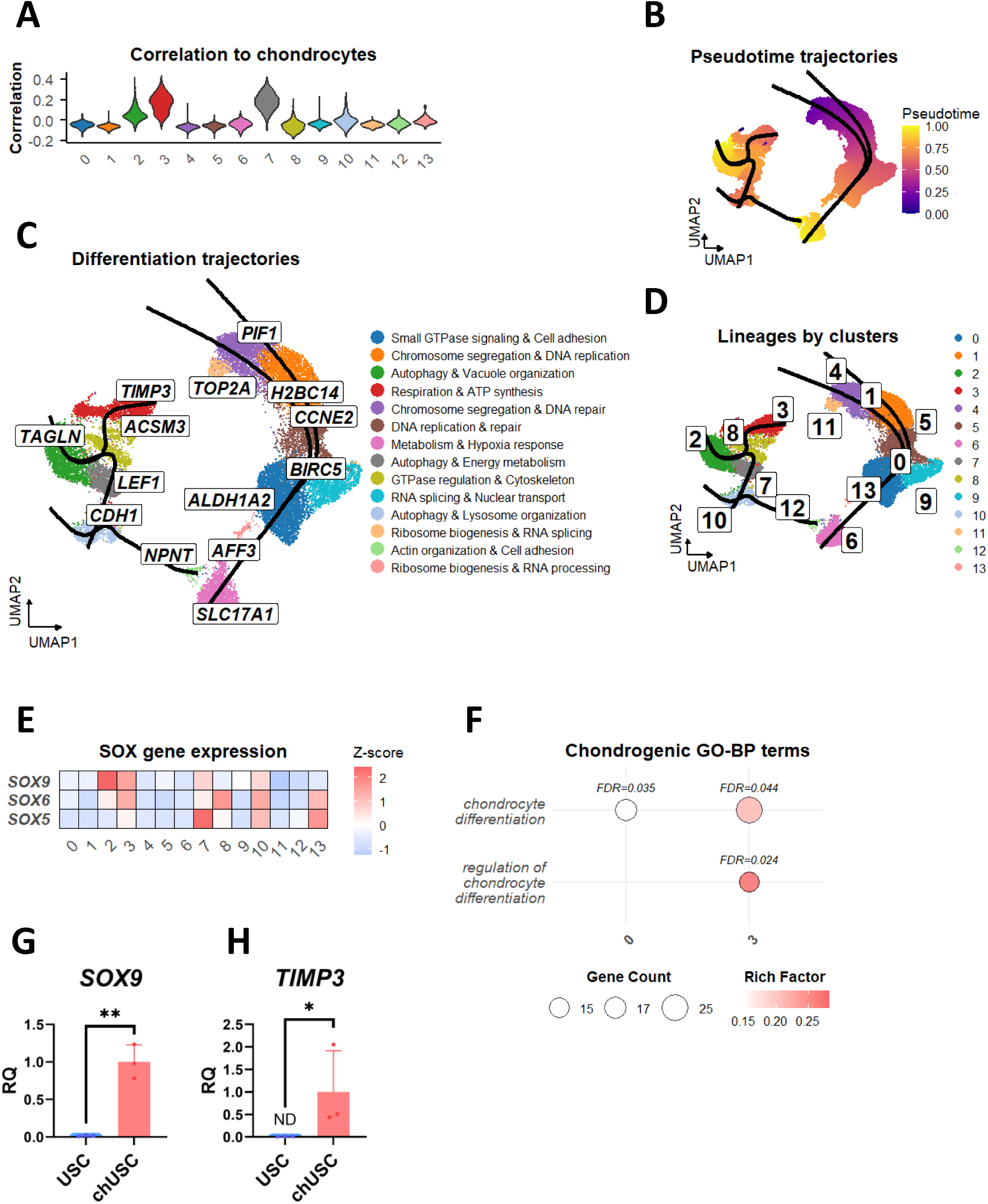
Single-cell lineage trajectories toward chondrocyte-like states. A: Violinplot highlighting correlation to chondrocyte reference cluster across merged undifferentiated and chondrogenic USC object clusters. (Cluster 3 pearson R = 0.164 ± 0.092, n = 2753). B: UMAP of merged USC object with slingshot pseudotime gradient and chondrogenic lineage curves. C: UMAP of the merged USC object with top marker annotation, chondrogenic lineage curves and summary of biological process GO-terms. D: UMAP of the merged USC object with cluster number annotation E: Heatmap, depicting the z-score normalized expression of SOX-transcription factors among clusters. F: Dotplot depicting clusters, enriched for chondrocyte related GO-terms. G/H: *SOX9* (RQ undifferentiated = 0.017 ± 0.001, differentiated = 1.00 ± 0.23, n = 3, p < 0.01, Welch’s t-test) and *TIMP3* (RQ undifferentiated = 0.011 ± 0.013, differentiated = 1.00 ± 0.91, n = 3, p < 0.05, Welch’s t-test) expression levels in undifferentiated and chondrogenic cells as measured by RT-qPCR in chondrogenic cells and undifferentiated controls. Bar with undetermined CT-values or exceeding 40 cycles, was highlighted (=ND).

To better understand how USCs transition from an undifferentiated to a chondrogenic state, we next analyzed how gene expression evolves along the trajectory. Pseudo time analysis using Slingshot revealed distinct differentiation trajectories emerging from proliferative *TOP2A*⁺ stem-like cells and progressing towards chondrocyte-like end states (Fig. 3B). This allowed us to pinpoint which genes become activated or silenced as cells progress toward a chondrocyte-like phenotype. We therefore looked for genes, differentially expressed in individual clusters along pseudo time. In the middle of our pseudo time, a subpopulation upregulated the expression of *ALDH1A2*, a gene involved in retinoic acid signaling. This means that retinoic acid is involved in chondrogenic modulation of USCs, highlighting the importance of vitamin metabolites in the differentiation process. At the end of the trajectory, we identified two distinct group of cells characterized by high expression of *TIMP3*, or *CDH1* (Fig. 3C). *TIMP3* is associated with extracellular matrix organization, whereas *CDH1* facilitates cell adhesion. Both of these features indicate that the cells are acquiring structural and regulatory functions typical of mature cartilage tissue, underscoring that USCs differentiation process recapitulates bona fide chondrogenesis.

To further strengthen these findings, we performed a Gene Ontology (GO) enrichment analysis and summarized the top 10 terms per cluster (Fig. 3C). Late-stage clusters with chondrocyte-like features (clusters 2, 3, 7) showed enrichment for processes related to ECM remodeling, cytoskeletal organization, and metabolic adaptation, including autophagy and energy metabolism pathways. These results again highlight the upregulation of *TIMP3*, together with *TAGLN*, as key markers of a matrix-producing, contractile phenotype emerging during chondrogenic maturation.

To assess if *bona fide* chondrogenic biological processes are active in these cells, we specifically looked for chondrocyte specific GO-terms. Here, the *ALDH1A2*⁺ intermediate was enriched for the GO-term “chondrocyte differentiation”. Again, suggesting a crucial function of retinoic acid related chondrogenesis modulation in USCs. Cluster 3, which contains most of aforementioned *TIMP3*⁺ cells, showed even stronger enrichment for the term “chondrocyte differentiation”, in addition to “regulation of chondrocyte differentiation” (Fig. 3F), supporting resemblance of chondrocyte transcriptional programs in chondrogenic USCs. Importantly, when we compared each cluster to a reference atlas of native chondrocytes, cluster 3 exhibited the highest transcriptional similarity. This confirms that these late-stage *TIMP3*⁺ cells represent the terminally differentiated end point of the USC chondrogenic trajectory, mimicking transcriptional gene expression programs active in chondrocytes.

We next examined the expression of the chondrogenic transcription factors *SOX9*, *SOX5*, and *SOX6*, known as the core regulatory trio of cartilage formation, to investigate the underlying transcription factor activity driving chondrogenesis in USCs. These genes were predominantly expressed in late clusters along pseudo time (Fig. 3B, D, E), confirming that USC-derived chondrocyte-like cells activate the canonical transcriptional network that drives cartilage lineage commitment. This transcriptional alignment with native chondrocytes supports the authenticity of the differentiation process.

To validate the single-cell findings at the bulk level, we quantified by RT-qPCR the expression of genes that resemble USCs bona fide chondrogenesis best. We chose *TIMP3*, as its expression was associated with chondrocyte related GO-terms (cluster 3, Fig. 3F) and chondrocyte similarity (Fig. 3A), and *SOX9*, as its expression increased in concordance with pseudo time (Fig. 3B, D, E). Both *SOX9* and *TIMP3* were strongly upregulated after differentiation (Fig. 3G, H), confirming the transcriptional activation observed in cluster 3. *TIMP3*, in particular, is an extracellular matrix regulator crucial for cartilage maintenance, further highlighting the functional maturation of these cells, driven by the chondrogenic master transcription factor *SOX9*.

Together, these results show that USC differentiation proceeds through defined transcriptional stages, from proliferative progenitors to mature chondrocyte-like cells. This is accompanied by activation of canonical cartilage transcription factors and functional pathways. The emergence of a *TIMP3*⁺ cluster with strong chondrocyte similarity and chondrogenesis related gene expression profile demonstrates that USCs can effectively recapitulate the molecular program of endochondral chondrogenesis.

### Potential for clinical application of USCs

To assess the clinical suitability of urine-derived stem cells (USCs), we established a xeno-free culture protocol by replacing fetal bovine serum (FBS) with autologous human serum (HS) during isolation and expansion (Supplementary Fig. 3A, B). This modification eliminates animal-derived components, a critical requirement for translational use in regenerative medicine.

When we compared transcriptional markers between FBS- and HS-expanded USCs, HS cultures showed increased expression of *MYC* (Fig. 4A, B). Elevated *MYC* is associated with proliferative, stem-like activity, whereas *KLF4* supports maintenance of a parietal epithelial cell (PEC)-like identity (as shown in Fig. 1D–I). Together, these findings indicate that HS expansion sustains the native USC phenotype, retaining both stemness and renal lineage features. Maintaining this balanced phenotype is desirable because it preserves the cells’ proliferative capacity and differentiation competence, both essential for efficient chondrogenic induction and clinical scalability.

**Figure 4:**
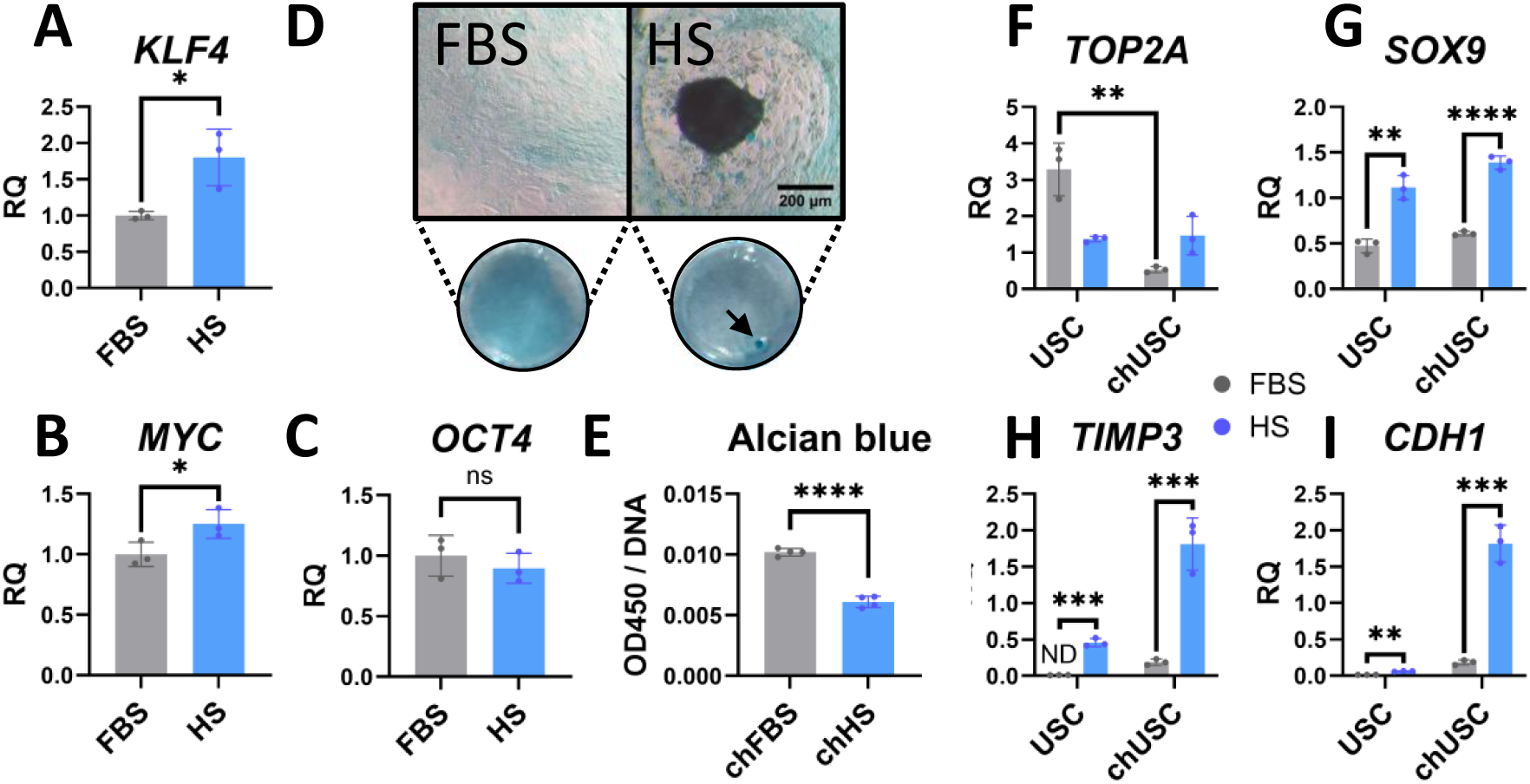
Xeno-free (HS) expansion enhances clinically relevant chondrogenesis. A/B: *KLF4* (RQ FBS = 1.00 ± 0.06, RQ HS = 1.80 ± 0.39, n = 3, p < 0.05, Welch’s t-test), *MYC (*RQ FBS = 1.00 ± 0.11, RQ HS = 1.25 ± 0.12, n = 3, p < 0.05, Welch’s t-test) and *OCT4* (RQ: FBS = 1.00 ± 0.17, HS = 0.90 ± 0.12, n = 3, p = 0.44, Welch t-test) expression levels in FBS-USC (FBS) and HS-USC (HS) cells as measured by RT-qPCR C: Alcian blue quantification of chondrogenic FBS-USC (chFBS) or chondrogenic HS-USC (chHS). D: Alcian blue staining of chFBS or chHS (OD650/DNA in FBS = 0.01 ± 0.0003, OD650/DNA in HS = 0.006 ± 0.0004, n = 4; p < 0.0001, unpaired t-test). E/F/G/H: RQ values of *TOP2A* (FBS: RQ undifferentiated = 3.28 ± 0.72; RQ differentiated = 0.54 ± 0.08, n = 3, p < 0.001, Welch’s t-test), *SOX9* (RQ chFBS = 0.61 ± 0.02; RQ chHS = 1.39 ± 0.07; n = 3; p < 0.0001, Welch’s t-test), *TIMP3* (RQ chFBS = 0.19 ± 0.04; RQ chHS = 1.81 ± 0.36; n = 3; p < 0.001, Welch’s t-test) and *CDH1* (RQ chFBS = 0.18 ± 0.03; RQ chHS = 1.81 ± 0.25; n = 3; p < 0.001, Welch’s t-test) of uninduced USCs (USC) and chondrogenic (chUSC) cells in pellet culture, comparing FBS or HS culture conditions. Bar with undetermined CT-values or exceeding 40 cycles, was marked (=ND).

To ensure that HS expansion does not trigger reprogramming toward an undesired pluripotent state, we measured *OCT4*, a canonical pluripotency transcription factor that can become aberrantly reactivated in dedifferentiating renal cells. *OCT4* expression remained unchanged (Fig. 4C) between FBS- and HS-expanded USCs. Additionally, proliferation appeared to be slightly reduced (Supplementary Fig. 3C), supporting that cells were not undergoing malignant transformation. Taken together, these data show that HS conditions preserve differentiation competence without inducing pluripotency.

We next evaluated whether xeno-free expansion affects the chondrogenic potential of USCs by inducing differentiation in monolayer culture, to observe differences in cellular morphology in both culture conditions. HS-expanded USCs (chHS) formed dense multicellular condensates, characteristic of mesenchymal condensation during early cartilage development, whereas FBS-expanded controls (chFBS) exhibited weaker condensation (Fig. 4D). We then quantified GAG content to observe differences in matrix deposition. FBS-expanded cells accumulated more GAGs (Fig. 4E), suggesting that FBS drives matrix deposition, whereas HS supports the initial cellular organization required for lineage commitment.

Because three-dimensional culture more closely mimics in vivo cartilage formation, we performed pellet differentiation of HS- and FBS-expanded cells. Under HS conditions, expression of *TOP2A*, the gene upregulated in proliferative progenitors, remained stable following induction, whereas it was markedly reduced under FBS (Fig. 4F). This indicates that HS preserves a pool of proliferative progenitors during early chondrogenesis, while FBS drives premature cell-cycle exit.

We then moved on validating the expression of genes appearing to reflect USCs chondrogenic program in our single cell atlas (Fig. 3), to see if xeno-free cultivation improves functional chondrocyte maturation in USCs. As differentiation progressed, HS pellets exhibited higher expression of the master chondrogenic regulator *SOX9* (Fig. 4G) and markedly increased levels of terminal associated genes *TIMP3* and *CDH1* (Fig. 4H, I). The upregulated expression of *TIMP3* corresponds to the late pseudo time chondrocyte-similar (Fig. 3A) and chondrocyte related GO-term related state (Fig. 3F) identified in our single-cell analysis, confirming that HS expansion promotes robust and complete chondrogenic differentiation. The increased *CDH1* expression, a gene involved in cellular aggregation, in 3D is in line with the observed chondrogenic aggregates in 2D (Fig. 4D), delivering orthogonal evidence that FBS diminishes chondrogenic outcomes of USC differentiation.

Together, these results demonstrate that autologous human serum supports xeno-free USC expansion while maintaining a proliferative, *MYC*-positive progenitor pool. During differentiation, HS-expanded cells efficiently activate *SOX9*-driven chondrogenic programs and progress toward *TIMP3*⁺/*CDH1*⁺ terminal states, indicating faithful recapitulation of cartilage maturation. Thus, making xeno-free cultured USCs an ideal starting material for chondrocyte regeneration approaches.

## Discussion

Our goal was to establish a non-invasive, physiologically relevant, and translational model of human cartilage development by leveraging urine-derived stem cells (USCs). By integrating high-resolution single-cell transcriptomic profiling, functional lineage differentiation assays, and a xeno-free expansion protocol, we aimed to delineate the cellular hierarchy and transcriptional programs underlying USC chondrogenesis, evaluate their potential to recapitulate native cartilage formation, and develop a clinically compatible platform for modeling genetic skeletal disorders and advancing personalized regenerative therapies.

To establish USCs as a well-defined biological system, it was necessary to resolve their cellular heterogeneity and origin. We began by deconstructing the cellular composition of bulk urine-derived stem cells to define cellular diversity. Single-cell RNA sequencing of USCs revealed nine transcriptionally distinct clusters, delineating a structured progenitor hierarchy rather than a uniform population (Fig. 1A, B). To definitively confirm their source within the kidney, we mapped our data against a reference atlas, which confirmed their parietal epithelial cell (PEC) origin (Fig. 1C-F), reinforcing the concept that USCs retain a tissue-specific transcriptional memory after ex vivo expansion, a process in line with behavior of adult stem cells^38^. This tissue-specific memory is beneficial as it provides a stable molecular identity that may preserve lineage fidelity during differentiation, enhancing the reproducibility of experiments and increasing their relevance for modeling kidney- or cartilage-related pathologies. Furthermore, defining both the heterogeneity and origin of USCs establishes a foundation for rational selection and manipulation of specific subpopulations to optimize chondrogenic or osteogenic differentiation for translational applications.

Next, we sought to understand the regulatory drivers behind the proliferative and quiescent compartments we observed. CytoTRACE and VIPER analyses identified *MYC* and *E2F4* as central regulators of the proliferative compartment (Fig. 1G-I). While MYC is classically associated with oncogenesis^39^, here it appears to drive physiologically controlled cell-cycle re-entry, supporting expansion without evidence of transformation^40^. Interestingly, although *E2F4* is often viewed as a repressor in classical cell-cycle control^41^, in certain developmental or stem-cell contexts it can act as a transcriptional activator. For example, in mouse embryonic stem cells, *E2F4* promotes expansion by directly activating cell cycle genes independently of the RB family, and E2F4 knockout slows S-phase entry and reduces viability^42^. This is particularly novel because *E2F4* has not previously been implicated in maintaining controlled proliferative activity in adult, non-pluripotent stem cell populations and its activation in USCs highlights a previously unrecognized mechanism that supports stable expansion. *KLF4* activity marked the quiescent PEC-like clusters (Fig. 1E-I), reflecting a retention of renal lineage traits, suggesting that these cells maintain a baseline functional identity even under proliferative stress.

This balance between proliferative and quiescent states is beneficial because it preserves both the stem-like potential and tissue-specific memory of USCs, enabling predictable differentiation into osteogenic and chondrogenic lineages while maintaining a physiologically relevant transcriptional profile. By identifying the molecular drivers of these distinct compartments, we gain insight into how USCs can be selectively manipulated to enhance expansion, lineage commitment, and ultimately, translational utility. Collectively, these data define USCs as a hierarchically organized progenitor population with proliferative and quiescent, lineage-primed cells. This architecture provides the mechanistic transcription factor activity driving their observed multipotency and establishes a foundation for subsequent chondrogenic studies.

Having defined their progenitor identity, we moved beyond transcriptional profiles and functionally validate their capacity to generate the skeletal lineages relevant to our model, osteogenic and chondrogenic fates. Complementary bulk RNA-Seq and RT-qPCR analyses showed upregulation of canonical osteogenic and chondrogenic markers (Fig. 2C–F), demonstrating that USCs can faithfully execute mesodermal lineage-specific programs. This functional validation is important because transcriptional similarity alone does not guarantee actual differentiation potential. Confirming that USCs can produce mature lineage-specific phenotypes strengthens their relevance as a cellular model. Moreover, this predisposition aligns with their renal epithelial origin, which may prime them for mesenchymal-like transitions under appropriate cues. Parietal epithelial cells, the presumed source of USCs, have been shown to undergo epithelial-to-mesenchymal transition and adopt progenitor-like phenotypes in vitro and in vivo^43,44^, suggesting that these cells harbor latent plasticity that may facilitate skeletal lineage reprogramming in our differentiation assays. This context-dependent flexibility likely underlies the ability of USCs to efficiently adopt osteogenic and chondrogenic fates, bridging epithelial and mesenchymal lineages. This indicates that USCs are not only transcriptionally competent but also functionally poised for skeletal differentiation, increasing their reliability for modeling cartilage and bone development, studying disease mechanisms, and ultimately serving as a scalable and patient-specific platform for regenerative applications.

As our results showed a marked bias toward chondrogenesis, we employed complementary 2D and 3D culture systems because they provide different but essential insights. Monolayer cultures allowed analysis of gene expression and cellular morphology during differentiation, while 3D spheroid cultures recapitulated key morphogenetic events such as mesenchymal condensation and extracellular matrix (ECM) deposition. To confirm that the matrix produced in 3D was authentic cartilage, we performed staining for the cartilage-specific proteoglycan aggrecan, a definitive marker of articular cartilage extracellular matrix structure and function^45^. The strong aggrecan deposition observed validates that our 3D differentiated tissues replicate key compositional features of native cartilage, thereby supporting their relevance for developmental and regenerative applications (Fig. 2G). This identifies USCs as a powerful tool for studying cartilage biology. Their intrinsic preference toward chondrogenesis is particularly attractive, as it indicates that these cells naturally favor to recreate cartilage-specific microenvironments, enabling more reliable modeling of cartilage development, disease processes, and potentially regenerative therapies.

While bulk assays confirmed differentiation potential, we aimed to unravel the precise sequence of cellular events and identify key transitional states during chondrogenesis at single-cell resolution. We therefore performed pseudo time analysis (Fig. 3B), which revealed the chondrogenic trajectory emanating from *TOP2A*⁺ progenitors, converging through an *ALDH1A2*⁺ intermediate enriched for chondrocyte-specific programs (Fig. 3C–F). The presence of this intermediate in undifferentiated populations suggests pre-patterning at the metabolic level via retinoic acid signaling^46^, highlighting the nuanced interplay between cellular state and lineage bias. This finding uncovers early regulatory checkpoints that may prime cells for chondrogenic commitment, providing mechanistic insight into how lineage bias is established. By defining these transitional states, we gain the ability to predict, manipulate, or enhance chondrogenic differentiation, thereby improving the fidelity of USCs as a model for cartilage development and disease, and informing strategies for regenerative therapies.

A critical question was whether the endpoint of our chondrogenic differentiation resembled genuine chondrocytes. Terminal differentiation yielded discrete chondrocyte-like subpopulations: a *TIMP3*⁺, and a *CDH1*⁺ cluster (Fig. 3A-D). TIMP3, a tissue inhibitor of metalloproteinases abundantly expressed in mature cartilage^47^, plays a crucial role in maintaining extracellular matrix integrity and protecting against proteolytic degradation^48^. Its strong induction therefore signifies the acquisition of a regulatory, matrix-stabilizing phenotype characteristic of functional chondrocytes. In parallel, the emergence of CDH1⁺ cells reflects activation of intercellular adhesion mechanisms. E-cadherin (*CDH1*) has been shown to enhance chondrogenic differentiation in human mesenchymal stem cell aggregates, acting together with N-cadherin to promote mesenchymal condensation and lineage commitment^49^. This underscores the importance of *CDH1* in facilitating mesenchymal condensation, an early and indispensable step of chondrogenic differentiation. Together, the coordinated expression of these structural and regulatory markers indicates that USCs undergo a structured, stepwise maturation process akin to endochondral development. This single-cell resolution map reveals not only the hierarchical differentiation of USCs but also actionable molecular targets, such as *TIMP3*-mediated matrix regulation and *CDH1*-associated adhesion, which may be leveraged to modulate differentiation or tissue formation.

Since a primary advantage of USCs is their non-invasive nature, we engineered a xeno-free protocol to make the entire procedure clinically relevant. We first asked whether xeno-free expansion alters the fundamental identity of USCs. HS-expanded USCs modestly increased *MYC* and *KLF4* expression compared with FBS, while *OCT4* remained unchanged, confirming safety with respect to pluripotency activation (Fig. 4A–C). This coordinated, increased, expression of stemness-associated and renal lineage-associated transcription factors is especially advantageous because it allows USCs to expand efficiently while retaining their original renal identity, minimizing the risk of dedifferentiation or acquisition of unintended phenotypes. This makes HS-expanded USCs a safer and more reliable starting material for translational applications, including patient-specific disease modeling and regenerative therapies.

We then rigorously compared the differentiation efficacy of HS-versus FBS-expanded cells to ensure that the clinical suitability did not compromise functional outcomes. During 2D differentiation, HS-expanded cells formed pronounced mesenchymal condensates, indicative of bona fide chondrogenesis^37^, whereas FBS-expanded cells accumulated more bulk glycosaminoglycans (Fig. 4D–E), reflecting accelerated ECM deposition. This comparison is beneficial because it highlights that xeno-free conditions do not simply mimic conventional culture but instead preserve a more physiologically relevant developmental program. To resolve this apparent discrepancy and better approximate in vivo cartilage formation, we switched to 3D pellet differentiation and gene expression analysis. In this context, HS-expanded USCs maintained *TOP2A* expression while robustly activating chondrogenic programs, corresponding to the late pseudotime chondrocyte-like states identified in our single-cell analyses (Fig. 4F-I). This finding is particularly advantageous because it demonstrates that xeno-free expansion strikes an optimal balance, by retaining a pool of proliferative progenitors necessary for long-term differentiation capacity while allowing faithful progression toward mature, functional chondrocytes. This balance ensures that USCs can serve as a reproducible, clinically compatible platform for cartilage modeling and regenerative applications, combining safety, scalability, and developmental fidelity.

To conclude, our study successfully establishes urine-derived stem cells (USCs) as a non-invasive, physiologically relevant, and translational model of human cartilage development. By combining single-cell transcriptomic mapping with functional differentiation in 2D, spheroid, and pellet cultures under xeno-free conditions, we delineated a hierarchical structure of USC chondrogenesis, from proliferative *MYC*/*E2F4*-active *TOP2A*⁺ progenitors through *ALDH1A2*⁺ intermediates to terminal *TIMP3*⁺ chondrocyte-like cells. This framework not only recapitulates key stages of native cartilage formation but also confirms that USCs preserve both lineage fidelity and differentiation potential during expansion. Together, these findings demonstrate that USCs provide a robust, clinically compatible platform for modeling genetic skeletal disorders and advancing personalized regenerative therapies, effectively bridging mechanistic insight with translational application.

## Methods

### Urine stem cell isolation and culture

USCs were isolated and expanded by modifying an existing protocol from Culenova et al ^50^. Briefly, urine was centrifuged at 300 g, cell pellet was washed with PBS 1% Penicillin Streptomycin (Gibco) and pelleted cells cultured after repeated centrifugation. Culture medium consisted of a 1:1 mixture of DMEM high glucose (Gibco), containing 1 % NEAA (Gibco) and 1 % Penicilin Streptomycin (Gibco), and KSFM (Gibco) with addition of 15 % FBS (Sigma Aldrich or Roth) and 5 ng/mL bFGF (Peprotech) or humankine thermostable bFGF (Sigma). For an animal free workflow, we harvested USCs in the same manner, this time resupending the cellular pellet in either Alpha MEM Eagle medium (PAN Biotech) supplemented 5 ng/mL fibroblast growth factor 2 (FGF2, Sino Biological) with 10% FBS (Sigma-Aldrich or Roth) as the FBS-Control, or in 10 % autologous human serum. All cell lines generated for this study were from the same donor, and no passages greater 5 were used. Culture medium was exchanged three times per week.

### Proliferation curves

To compare proliferation, we seeded FBS- and HS-derived USCs of the same passages in triplicates onto 96 well plates at a density of 1000 cells per well, and counted cells manually with trypan blue exclusion every second day. Doubling time was calculated for cells in the exponential growth phase (day 4-8) as the following:

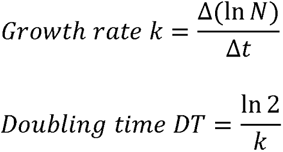

### Urine stem cell differentiation and staining

Chondrogenic differentiation was induced with the StemPro Chondrogenesis Kit (Gibco). Osteogenic induction was facilitated with an osteogenic medium described by zhang et al ^51^. Differentiation lasted for two weeks with media change three times per week. To stain glycosaminoglycans in chondrogenic, and calcium deposits in osteogenic cells, we fixed cells with 4 % paraformaldehyde, washed with PBS, and stained for one hour with alcian blue or alizarin red s staining solution. After a final washing step with PBS images were taken under a VertA1 (Zeiss) microscope. A separate USC line, harvested at a different time point from the same donor, was cultured for 2 days in Alpha MEM Eagle medium (PAN Biotech) supplemented with 10% fetal bovine serum (FBS, Sigma-Aldrich or Roth) and 5 ng/mL fibroblast growth factor 2 (FGF2, Sino Biological), and served as the non-induced control.

For a comparative pellet culture between HS- or FBS-USCs, we centrifuged 250000 USCs at 300 g for 10 min in 15 ml falcon tubes (Corning) and incubated with in house made chondrogenic media. The media recipe was: DMEM HG with pyruvate (Gibco), human recombinant insulin 6.25 μg/mL (Merck), human holo-transferrin 6.25 μg/mL (Sigma Aldrich), sodium selenite 6.7 ng/mL (Sigma Aldrich), 10 % Serum (autologous human serum or FBS, Roth) 10 ng/mL recombinant human TGF-Beta 1 (Peprotech), 100 ng/mL recombinant human IGF1, 100 nm Dexmethason (Sigma Aldrich), 50 µg/mL ascorbate-6-phosphate (Sigma Aldrich).

Alcian blue quantification was performed by lysing cells for 2 hours at room temperature with 6M Guanidine hydrochloride (Sigma). Absorbance was measured at 650 nm. For normalization to DNA content, DNA was extracted from control wells with the DNeasy Blood & Tissue Kit (Qiagen).

### Single-cell RNA sequencing and data Processing

USCs and chondrogenic USCs were subjected to single-cell RNA sequencing (scRNA-seq) to characterize transcriptomic profiles in undifferentiated and chondrogenically differentiated states. Cell fixation, barcoding, cDNA amplification and Libraries were prepared using Parse Biosciences Parse Evercode WT V3 Workflow and sequenced on Illumina NovaSeq X Plus. FASTQ files were processed using Trailmaker^TM^ pipeline module (https://app.trailmaker.parsebiosciences.com/, pipeline v1.5.0, Parse Biosciences). Raw count matrices and associated metadata were processed in Seurat (version 5.3.0)^32^ in R. Cells with <1ST or >99th percentile of gene counts or Unique Molacular Identifier (UMI) counts, or >95th percentile mitochondrial reads, were excluded. Data were normalized (LogNormalize), variable features (n = 2,000) identified (vst), and counts scaled while regressing out nCount_RNA and percent.mt. PCA (dims 1–20) was followed by UMAP visualization and Louvain clustering (resolution = 0.5). Cluster markers were identified via Wilcoxon testing (min.pct = 0.25). For visualization of data we used Seurat’s DotPlot, VlnPlot or DimPlots and the R packages ggplot2 (version 3.5.2)^52^, ggrepel (version 0.9.6)^53^, viridis (version 0.6.5)^54^ and RColorBrewer (version 1.1.3)^55^.

### Tissue origin and cell-type annotation

Marker lists for human urinary and renal tissues were curated from CellMarker 2.0^33^, filtered for uniqueness across tissues, and scored per cell using Seurat’s AddModuleScore. Complementary AUC enrichment was computed with AUCell (version 1.30.1)^56^ on the normalized data. To assign putative cell identities, we leveraged a well-annotated human kidney atlas (Tisch et al.^34^) as a reference. Briefly, the atlas and our USC Seurat object were first normalized and dimensionally reduced in parallel. We then computed integration anchors between the reference and query using FindTransferAnchors. Cell-type labels stored in the reference metadata were projected onto the query via TransferData. Finally, we summarized mapping confidence by computing the median prediction.score.max within each Seurat cluster to guide downstream cluster ordering and interpretation.

### Stemness inference and transcription-factor activity mapping

CytoTRACE2 (version 1.1.0)^36^ was run on the top 10,000 variable genes (counts slot) to assign each cell a continuous stemness score. Cells were binned into five quantile-defined stemness groups. We inferred per-cell transcription-factor activities using VIPER (version 1.42.0)^57^ with high-confidence DoRothEA (version 1.20.0)^58^ A/B regulons on the Seurat-normalized data slot. TF activities were merged into the metadata alongside CytoTRACE2 scores and UMAP coordinates. For each stemness group (the five quintile bins), we computed the mean activity of every TF and selected the top three TFs uniquely highest in each group. To visualize regulatory dynamics, we computed Spearman correlations between TF activities and CytoTRACE2 scores across all cells, identifying the top 20 TFs most strongly associated with stemness.

### Cell cycle analysis

Cell cycle phase assignment was performed using the built-in Seurat gene sets for S phase and G2/M phase genes. The Seurat function CellCycleScoring was applied to the processed data, utilizing the predefined sets of cell cycle genes for the S and G2/M phases. Each cell was classified into one of the four phases: G1, S, G2/M, or unassigned. Phase information was then merged with cluster identities, and the proportion of cells in each cell cycle phase per cluster was calculated.

### Bulk RNA-sequencing

RNA was extracted from USC samples with RNeasy Mini Kit (Qiagen)with DNAse digestion, according to manufacturer. A cDNA library was generated for each sample using the Illumina stranded mRNA kit. Paired-end sequencing of the libraries with a fragment length of 159bp was conducted on an Illumina NovaSeq6000 platform. Raw data was then converted into reads and demultiplexed using Illumina Dragen Software (v. 3.8.4).

Unwanted RNA was filtered from the reads using bwa mem (version 0.7.17-r1188, arXiv:1303.3997) in combination with samtools (version 1.17^59^) and converted back into fastq format using the SamToFastq tool from GATK (version 4.2.1.0^60^). Reads from each sequencing lane were aligned individually to the hg38 reference genome with Ensembl (release 110^61^) gene annotations using STAR (version 2.7.10a^62^). For each sample, resulting alignments were combined using the MergeSamFiles command from Picard (version 2.25.4, http://broadinstitute.github.io/picard/). Gene quantification in form of a count matrix was generated using featureCounts (version 2.0.1^63^) and Ensembl gene annotations corresponding to those of the alignment reference. Alignment and quantification levels were used for quality assessment.

### Differential gene expression analysis

Raw gene-level count data were obtained from bulk RNA-sequencing of three undifferentiated and three differentiated (osteogenic or chondrogenic) USC samples. Differential gene expression (DGE) analysis was conducted using the DESeq2 package (version 1.48.1)^64^ in R (version 4.5). For each comparison, gene-level counts were used to construct a DESeqDataSet object, with group labels assigned as the experimental condition. The DESeq2 pipeline was run with default parameters using DESeq(), and differential expression results were extracted using the results() function, specifying contrasts such that log2 fold changes represented upregulation in the differentiated condition relative to WT. Genes with an adjusted p-value (Benjamini-Hochberg method) less than 0.05 were considered statistically significant. Gene annotations were retrieved by mapping Ensembl IDs to HGNC symbols using the org.Hs.eg.db package (version 3.21.0)^65^. Differential expression results were visualized using volcano plots created with the ggplot2^52^ and ggrepel^53^ packages. Sample metadata was generated to define the experimental condition associated with each sample (WT, chWT, oWT), and this metadata was used to create a DESeqDataSet object using the DESeq2 package. The dataset was then log-transformed using the regularized log (rlog) transformation to stabilize variance across the dynamic range of expression levels. To improve PCA interpretability and reduce noise, features (genes) with negligible or zero variance were removed. Columns (genes) with variance less than 1e-6 or equal to zero were filtered out. Rows (samples) with zero variance across retained genes were also removed. PCA was performed on the transposed, filtered, and normalized gene expression matrix using the base R prcomp() function with centering and scaling enabled. The first three principal components were extracted for visualization. The resulting PCA scores were visualized using the plotly package^66^ to generate an interactive 3D scatter plot. The final plot was exported as an HTML widget using the htmlwidgets package^67^.

### Reverse transcriptase quantitative PCR

RNA was extracted from USC samples with RNeasy Mini Kit (Qiagen) with DNAse digestion, according to manufacturer. Alternatively, we used NucleoSpin RNA Plus (Machery Nagel) with gRNA removal column. We made sure to use samples prepared with the same procdures for comperative anlysis. For cDNA Synthesis LunaScript RT SuperMix Kit (New Englang Biolabs) was used. RT-qPCR was performed with TaqMan Master-Mix and predesigned Assays (Applied Biosystems) on the Quantstudio 12K Flex Platform (Applied Biosystems). *RPLP0* served as housekeeping gene. Values were calculated with the delta delta CT (ddCT) method and four technical replicates. Statistics (unpaired welch’s t-test of ddCT values) and visualization were performed in R studio and Graphpad Prism (version 10). We always used three biological replicates per condition.

### 3D-culture and immunofluorescence

20000 USCs were seeded in each well of a 96 well Nunclon Sphera ultra low attachment cell culture plate (ThermoFisher Scientific) and centrifuged for 10 minutes at 300 g to enable scaffold free cellular self-aggregation. Cells were cultured for 3 weeks with StemPro Chondrogenesis Kit (Gibco). As an undifferentiated control we used USCs from the same donor (different batch), which were cultured 2 days in Alpha MEM Eagle (PAN Biotech) with 10 % FBS (Sigma Aldrich or Roth) and 5 ng/mL FGF2 (SinoBiological).

For immunofluorescent staining, spheres were fixed for one hour with 4 % paraformaldehyde (PFA) at room temperature. In the following, the spheroids were permeabilized with Triton X-100 for 30 minutes and then blocked with 1 % bovine serum albumin (Sigma-Aldrich) in PBS for 45 minutes. For primary antibody incubation anti-aggrecan (ab3778, abcam) was used in a concentration of 1:500 for one hour, followed by one hour of secondary anti-mouse antibody (AlexaFluor 488, Invitrogen) in a 1:500 dilution, with DAPI (1:1000, thermo fisher scientific). After washing with PBS spheroids were mounted with aqua polymount (Polysciences) on concave slides (Paul Marienfeld). Pictures were taken with the Axio Imager 2 with Apotome (Zeiss) and 25 Z-stacks. We used the same exposure times for differentiated and undifferentiated spheroids (DAPI: 10 ms, Aggrecan: 35 ms). The same image enhancements for each condition, including normalization to DAPI, were performed with FIJI^68^.

### Merging undifferentiated and chondrogenic USC datasets

Undifferentiated and chondrogenic USC Seurat objects were annotated with a stage metadata field (“undiff” vs. “chondrogenic”). The two objects were merged and reprocessed end-to-end (NormalizeData, FindVariableFeatures, ScaleData, RunPCA, RunUMAP, FindNeighbors, FindClusters) to generate a unified embedding for downstream trajectory and mapping analyses. Results comparing chondrogenic and undifferentiated samples were obtained from DESeq2 analysis (adjusted p-value < 0.01, log2 fold change > 1). For proving concordance with bulk RNA-Sequencing, we extracted significantly upregulated genes and calculated a gene module score in the merged Seurat object. The AddModuleScore function was applied to quantify the expression of this gene set across cells. Resulting module scores were visualized on a UMAP embedding using the FeaturePlot function (Supplementary Fig. 2D, E).

### Construction of healthy cartilage reference atlas and label transfer onto merged USC dataset

We assembled a multi-sample single-cell reference from publicly available human cartilage data^69^. For each of the three selected samples, raw count matrices, feature, and barcode files were imported into Seurat. Cells with < 200 genes, < 3 cells per gene, or > 10% mitochondrial reads were excluded. Each sample was SCTransformed (regressing out percent.mt), then merged into a single Seurat object. We first applied the same QC filters as for our USC objects to the published cartilage reference to generate a high-quality “healthy cartilage” atlas. Cells annotated as normal cartilage were subsetted, and all assay layers were collapsed into a single RNA assay. This object was then log-normalized (NormalizeData), variable features were identified (FindVariableFeatures), data were scaled (ScaleData), and PCA (dims 1–30), neighbor-graph construction, clustering, and UMAP were performed. The merged undifferentiated□+□chondrogenic USC object underwent the same SCT-assay preprocessing. We then selected the top 3,000 integration features across both objects and computed label-transfer anchors with FindTransferAnchors (normalization.method = "SCT"). Finally, reference cluster identities were projected onto the USC dataset via TransferData, yielding per-cell predicted cluster labels and confidence scores, which were appended to the USC metadata (Supplementary Fig. 2F-H).

### Mapping and correlation of USCs to chondrogenic reference atlas and pseudotime inference

We loaded the processed human cartilage reference and the mapped USC dataset into Seurat (SCTransform normalization). To identify “cluster□4–like” USCs, we labeled any cell whose transferred reference cluster equaled “4” and whose PredictionScore□>□0.7 (Supplementary Fig. 2I). Next, we computed the mean SCT embedding for reference cluster□4 and, for each USC cell, calculated the Pearson correlation between its SCT profile and the cluster□4 embedding. These correlation coefficients were then overlaid on the merged UMAP embedding to show how closely individual USCs resemble cluster□4 in principal component space (Supplementary Fig. 2J, K). For pseudotime analysis, we ran Slingshot (version 2.16.0)^70^ on the UMAP data, using the mitotic (*TOP2A*⁺) cluster as the root. Slingshot automatically detected terminal states. We computed pseudotime values per cell, appended them to the USC metadata, and projected them onto the UMAP. Finally, we annotated clusters based on their top marker genes (FindAllMarkers) and performed Gene Ontology biological process enrichment (clusterProfiler, version 4.16.0)^71^. We mapped DE genes to Entrez IDs, ran enrichGO with Benjamini–Hochberg correction, and selected the top ten GO–BP terms per cluster. These process annotations, and their positions on the UMAP, were summarized and visualized using cluster centroids

### Chondrogenic cluster annotation and functional characterization

To identify clusters with chondrogenic potential, we integrated transcriptomic similarity to a reference chondrocyte population, GO enrichment, and expression of canonical marker genes. Average “cluster 4–like” scores (Pearson correlation to reference cluster 4) were computed per cluster and visualized with VlnPlot. The Z-score normalized, scaled average expression of chondrogenic markers (*SOX5*, *SOX6*, *SOX9*) were calculated and depicted in a heatmap. GO biological process enrichment was filtered to include only terms related to chondrocyte differentiation (GO:0032330, GO:0002062), and enrichment values were visualized with ggplot2 across clusters.

## Supporting information

Description of Additional Supplementary Files

## Data availability

All sequencing data generated in this study have been deposited in the Gene Expression Omnibus (GEO) database. scRNA-Seq of USCs: accession code GSE307232. Bulk-RNA-Seq of USCs: accession code GSE306729. RT-qPCR raw and analyzed tables, differential gene expression tables of bulk RNA-Seq confirming differentiation, Top 10 and chondrocyte related GO-Terms per cluster, as well as marker genes for clusters used in chondrogenic trajectory analysis are in Supplementary Data.

## Code availability

All the R codes used to generate the results of this study are publicly available on GitHub: https://github.com/alexschulzcell/USC-paper.

## Acknowledgments

We thank Mohammad Deen Hayatu and Evelyn Galsterer for technical assistance. This work was funded by the Deutsche Forschungsgemeinschaft (DFG, German Research Foundation) – [TH896 7-1].

## Author information

Contributions

A.S: performed the experiments, processed and analyzed the data, and wrote the original draft, E.B: processed the raw sequencing data. M.Z: revised figures and contributed to xenofree media development, A.S.B: revised manuscript, S.U: uploaded all sequencing data, A.B.E: assisted with sample collection, C.T.T: project P.I, acquired funding, supervised research and study design, and revised manuscript

## Ethics declarations

Competing interests

The authors declare no competing interests.

## Supplementary Information

## Notes

https://www.ncbi.nlm.nih.gov/geo/query/acc.cgi?acc=GSE306729

https://www.ncbi.nlm.nih.gov/geo/query/acc.cgi?acc=GSE307232

### Competing Interest Statement

The authors have declared no competing interest.

### Summary of Updates

Overall writing was significantly improved. OCT4 expression data (RT-qPCR) from FBS and HS cells was added.

