## Supplementary Information for "Single-cell dissection of cell hierarchies in urine-derived stem cells – application to chondrogenesis and general scalability"

**Supplementary Figures**

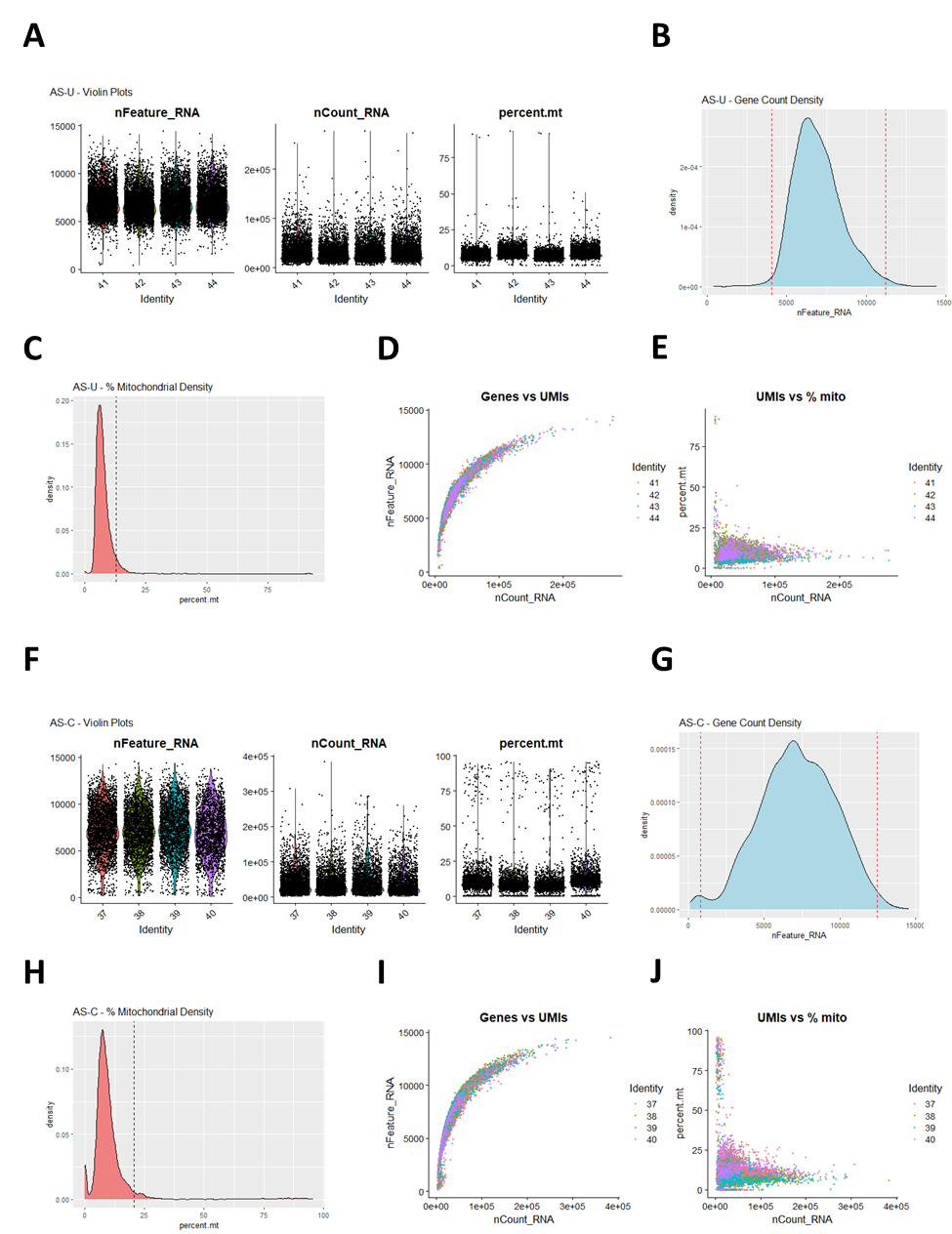

**Supplementary Figure 1: Quality control of single cell RNA-Sequencing data.** Panels A–E (AS U= undifferentiated) and F–J (AS C= chondrogenic) show the same set of QC visualizations for each condition. A/F: Violin plots of detected genes per cell (nFeature_RNA), UMI counts per cell (nCount_RNA), and percent mitochondrial reads (percent.mt). B/G. Density of detected genes with dashed lines marking the 1st and 99th percentile thresholds. C/H: Density of percent mitochondrial reads with a dashed line at the 95th percentile threshold. D/I: Scatter of UMI counts vs. detected genes, illustrating library complexity. E/J: Scatter of UMI counts vs. percent mitochondrial reads, highlighting high mt cells.

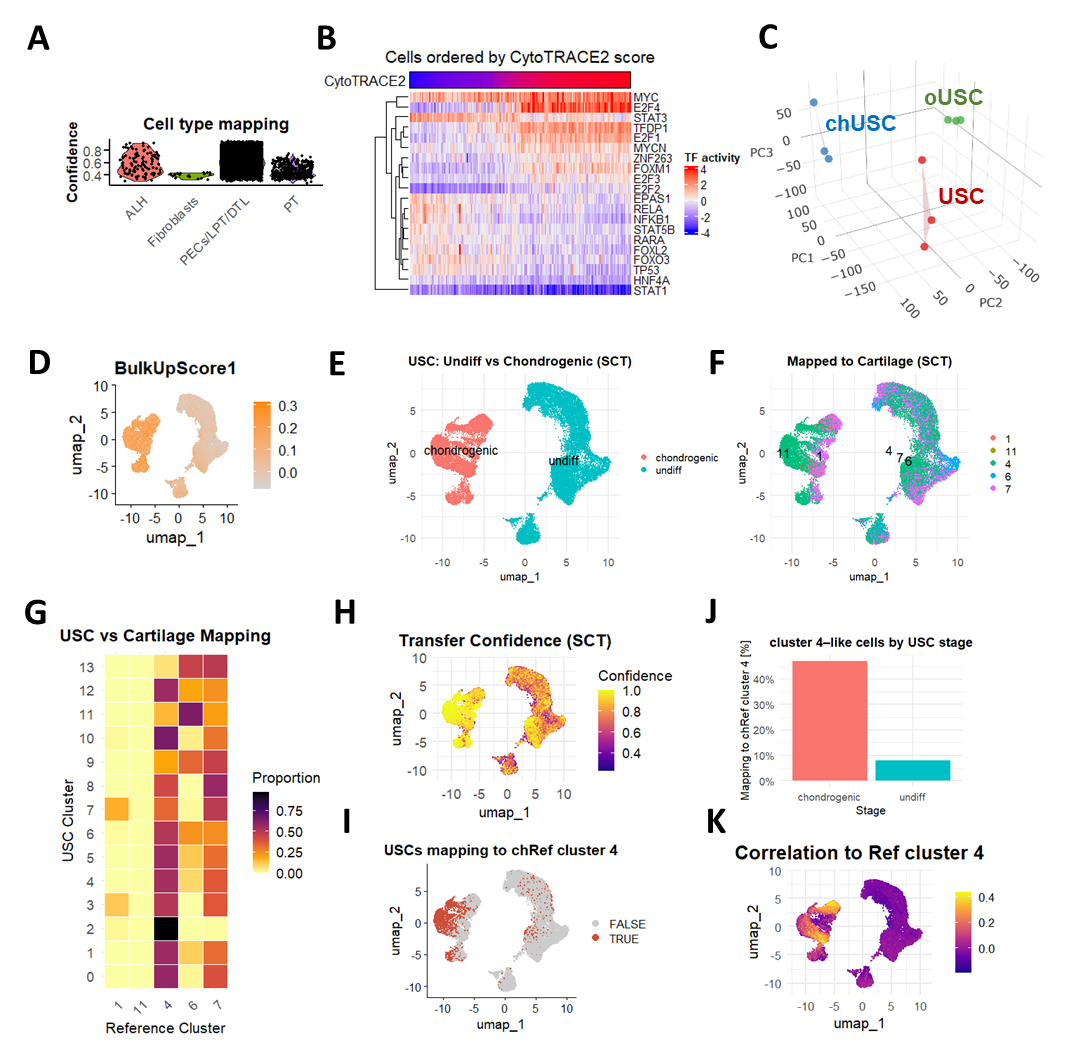

**Supplementary Figure 2: Supporting information for computational analysis.** A: Violinplot with predicted cell types for undifferentiated USCs. B: Top20 stemness related transcription factor heatmap. C: 3D-PCA of differentiated and undifferentiated replicates. D: BulkUpScore, reflecting upregulated genes in bulk RNA-Sequencing, highlighted in the merged undifferentiated and chondrogenic USC seurat object. E: Merged object UMAP with cell type annotations. F: Merged UMAP mapped to cartilage reference sample highlighting predicted reference labels. G: Heatmap visualizing mapping proportions of merged USC object to reference objects. H: UMAP of merged object with transfer confidence color gradient. I: UMAP of merged object, highlighting cells with mapping confidence to chondrocyte reference cluster > 0.7. J: fraction of cells mapping to the reference cluster across stages´. K: Correlation values with reference cluster mapped onto the merged UMAP.

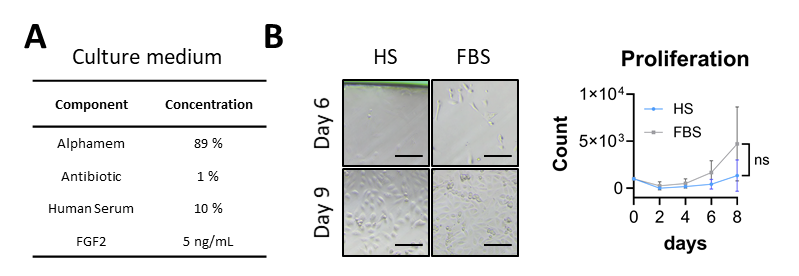

**Supplementary Figure 3: Expansion of USCs in HS conditions.** A: Medium recipe for USC expansion with autologous serum. B: Representative images of colonies emerging under HS or FBS expansion at day 6 and day 9. C: Proliferation curves of HS or FBS expanded USCs (n = 3).
