## Supplementary figures and images for "Single-cell dissection of cell hierarchies in urine-derived stem cells – application to chondrogenesis and general scalability"

### Supplementary Video

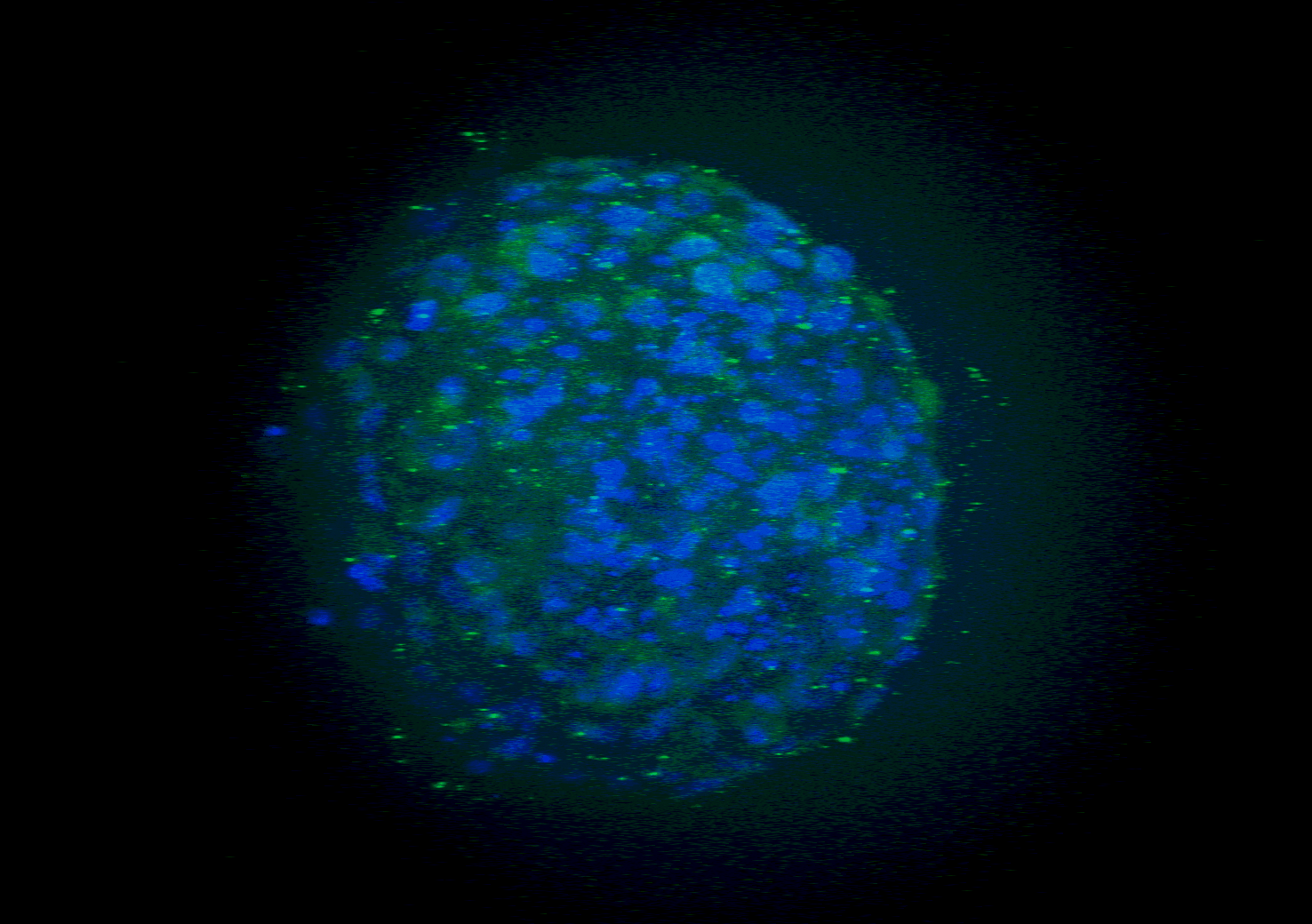
