## Supplementary material for "Single-cell dissection of cell hierarchies in urine-derived stem cells – application to chondrogenesis and general scalability": Description of Additional Supplementary Files

**Supplementary Data 1**

Contains the raw RT-qPCR tables.

**Supplementary Data 2**

Contains the analyzed RT-qPCR tables.

**Supplementary Data 3**

Contains the differential gene expression tables of bulk RNA-Seq of differentiation towards osteogenic/chondrogenic cells from DESeq2.

**Supplementary Data 4**

Contains tables of the top 10 and chondrocyte related GO-Terms per cluster of the chondrogenic trajectory analysis.

**Supplementary Data 5**

Contains the top marker genes for clusters used in chondrogenic trajectory analysis.

**Supplementary Video**

An animated Gif. file of a USC-derived chondrogenic spheroid.
